# Controlling Multicomponent Condensate Morphology via Additive-Modulated Interactions

**DOI:** 10.1101/2025.06.09.658693

**Authors:** Jiahui Wang, Arash Nikoubashman, Young C. Kim, Jeetain Mittal

## Abstract

The morphology of biomolecular condensates plays a critical role in regulating intracellular organization and function by enabling both spatial and temporal control over biochemical processes. Recent studies have shown that small-molecule cosolutes can not only modulate phase separation but also influence condensate morphology. However, the mechanistic understanding of how small molecules regulate condensate structure remains limited. In this study, we employ coarse-grained molecular dynamics simulations to investigate how the morphology of two-component condensates can be modulated through the introduction of additional small-molecule cosolutes. By systematically varying the interaction strengths between the small molecules and the macromolecular components, we observe morphological transitions between, e.g., core-shell and dewetted structures. To rationalize these transitions, we calculate second virial coefficients in the presence of the small molecules, providing a molecular-level framework to capture shifts in effective homotypic and heterotypic interactions. We further investigate the role of stoichiometry between the small molecules and macromolecules, demonstrating that stoichiometry and interaction strength jointly determine the condensate morphology by altering the relative interaction strengths among components. Additionally, we show that fully miscible two-component condensates can undergo transitions to microphase-separated morphologies, such as core-shell or dewetted, upon small molecule introduction. Together, these findings reveal that condensate morphology can be rationally tuned through interaction- and stoichiometry-dependent mechanisms, offering molecular-scale insights into how small-molecule cosolutes modulate condensate structure.

## Introduction

Macromolecular mixtures can undergo phase separation to form multiphase condensates, a phenomenon widely observed in biomolecular condensates^1–3^. The hierarchical organization of biomolecular condensates has been identified in various cellular structures^4–7^, such as the multiple phases in the nucleolus^6^ and the core-shell architecture in stress granules^8^, which are closely associated with condensate properties and functions^9^. For instance, within the nucleolus, distinct condensate phases organize ribosomal maturation steps in a sequential manner, maintaining structural separation and functional precision to optimize processing efficiency^6,10^. The ability of biomolecular condensates to form subcompartments enables fine-tuned positional and chronological regulation of complex intracellular processes, drawing increasing scientific attention.

Consequently, research on multiphase condensates has expanded in recent years, exploring their organizational mechanisms from various perspectives. Feric et al. demonstrated that the formation of multiple nucleolar subcompartments is driven by differences in surface tension^4^, while other studies have also highlighted the role of interfacial tension, an essential mesoscale property, in governing multiphase condensate formation^11–13^. By investigating the local microenvironments within biomolecular condensates, Ye et al. revealed that local micropolarity differences are crucial for the emergence of multiphase structures^14^. At the molecular level, Wells/Sojitra et al. proposed that the spatial segregation of condensates results from a balance between homotypic and heterotypic interactions^15^. Similarly, Rana et al. demonstrated that asymmetric oligomerization among condensate components regulates the miscibility of intrinsically disordered proteins (IDPs), further shaping the phase behavior of biomolecular condensates^16^.

Apart from understanding their formation mechanisms, recent efforts have also focused on how to regulate condensate morphology by modifying system composition, including the introduction of macromolecules^17–19^, crowding agents^20,21^, pH variation^22,23^, and salt concentration^24,25^. Notably, Zhu et al. reported that the introduction of small-molecule cosolutes, such as ions from the Hofmeister series, can alter condensate microenvironments and reshape condensate morphology^26^. These findings highlight the emerging importance of small-molecule cosolutes in modulating condensate architecture.

Indeed, small-molecule cosolutes have recently begun to be studied as modulators of biomolecular phase separation^27,28^. For example, Babinchak et al. showed that Bis-ANS, a fluorescent small molecule, acts as a biphasic modulator of protein condensation, capable of both inducing phase separation and preventing droplet formation^29^. ATP as a similar biphasic modulator has also been reported^30–32^. Wang et al. further demonstrated that small molecules such as Ro-3306 can alter the material properties of TFEB condensates, enhancing droplet size and fusion propensity, thereby modulating autophagosome-lysosome dynamics^33^. These studies highlight the important role of small-molecule cosolutes in biomolecular condensate research, providing powerful tools for uncovering the mechanisms of phase separation and offering promising strategies for therapeutic intervention in condensate-related diseases.

Despite growing evidence of the regulatory role of small molecules in biomolecular condensates, a fundamental understanding of how such small-molecule cosolutes influence the morphology of multicomponent condensates remains limited. Therefore, in this study, we aim to establish a general framework that directly links molecular interaction parameters to the mesoscale phase behavior of macromolecular systems, thereby elucidating the principles underlying morphology control through the introduction of small molecules. Recognizing that biomolecular condensates, typically composed of proteins and nucleic acids, often arise from multivalent interactions that can be effectively captured by simplified macromolecular models^2,15,34–36^, we employ simulations of a binary macromolecular system with the addition of small molecules. By systematically varying the interaction strengths between the small molecules and the macromolecular components, we investigate the resulting morphology transitions. This approach builds upon previous studies that have successfully utilized comparable models to probe diverse chemical phenomena^37–39^. For example, Li et al. employed a similar model to explore the fabrication of hybrid nanocolloids via the self-assembly of polymer blends and inorganic nanoparticles^39^. These studies collectively demonstrate that such generic models can serve as powerful tools for probing multicomponent interactions across diverse chemical contexts.

Our investigations reveal distinct morphological transformations, such as transitions from core-shell structures to dewetted states, which we quantitatively rationalize through theoretical calculations of the relative molecular interaction strengths. Furthermore, we explore the influence of stoichiometry on these morphological transitions, identifying a “re-wetting” phenomenon. Finally, we extend our analysis to examine the impact of small molecules on the morphology of miscible two-component condensates. Our findings offer a molecular-level perspective for understanding and potentially controlling the morphology of multicomponent condensate through the strategic incorporation of small-molecule cosolutes.

### Molecular Model and Simulation Methodology

We employed a coarse-grained polymer model, where each monomer is represented by a single bead, with a diameter *σ* = 5 Å and a mass of 100 g/mol, and solvent effects are treated implicitly through effective monomer-monomer pair interactions^40^(**Fig. 1a**). The system consists of two homopolymer species, chain A and chain B, each comprising 50 monomers of type A or B, respectively (*N* = 50). The small molecule was modeled as a single bead (type C), of the same size and mass as the polymer monomers.

**Fig. 1.**
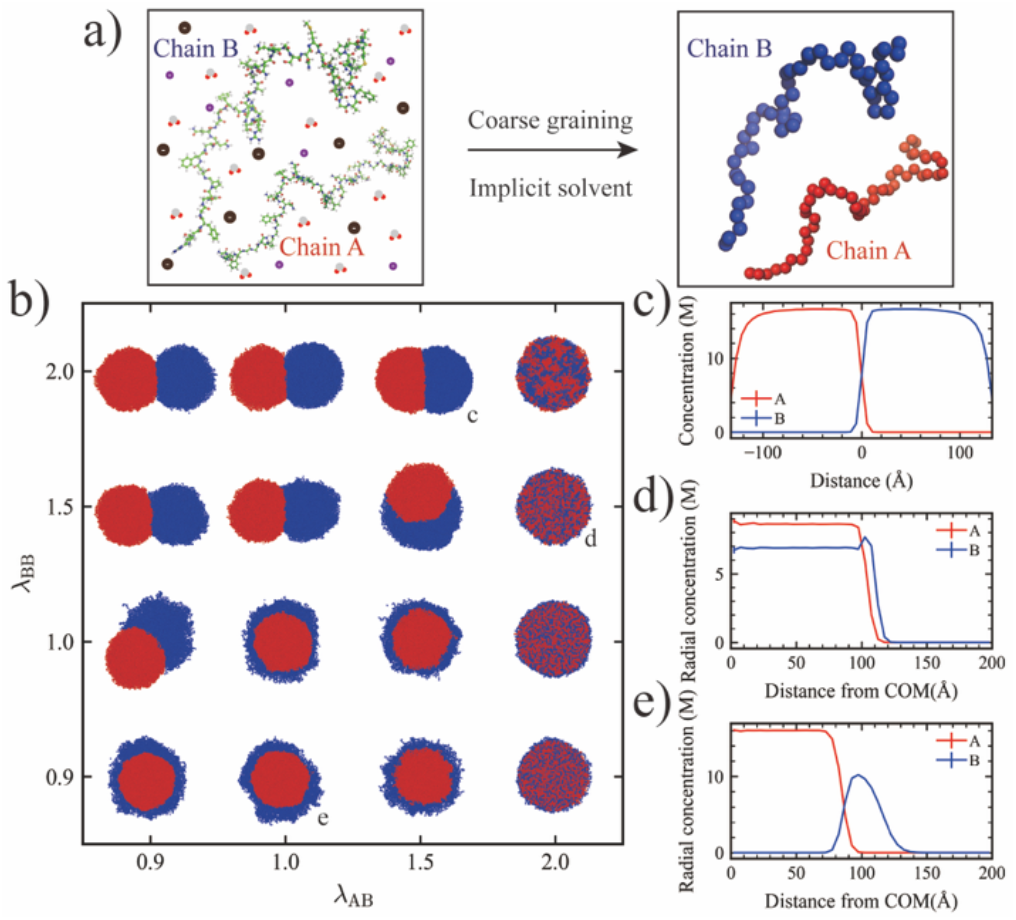
(a) Schematic of the implicit coarse-grained polymer model. (b) Morphological state diagram of two-component condensates at various combinations of *λ*_AB_ and *λ*_BB_, with fixed *λ*_AA_ = 2.0. Snapshot colors correspond to the component colors shown in the schematic. (c) Concentration profiles of A and B monomers along the axis connecting the centers of mass (COM) of the A-rich and B-rich regions (*λ*_AB_ = 1.5, *λ*_BB_ = 2.0). (d, e) Radial concentration profiles of A and B monomers measured from the COM of the entire condensate for (d) *λ*_AB_ = 2.0, *λ*_BB_ = 1.5 and (e) *λ*_AB_ = 1.0, *λ*_BB_ = 0.9.

Bonded interactions between adjacent monomers were modeled using a harmonic potential:

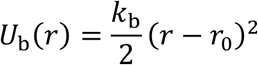

where *r* is the distance between bonded monomers, *k*_b_ = 20 kcal/(mol Å^2^: is the spring constant, and *r*_0_ = 3.8 Å is the equilibrium bond length.

Non-bonded interactions were described by a modified Lennard-Jones (LJ) potential, where the attractive contribution was scaled by a pairwise hydropathy parameter *λ*_ij_ between monomer types *i* and *j*:

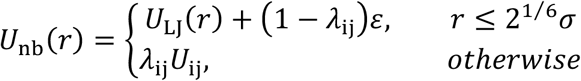

Here, *U*_LJ_(*r*) is the standard LJ potential:

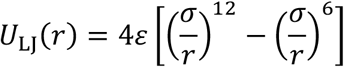

where *ε* = 0.2 kcal/mol is the interaction energy. By tuning the parameter *λ*_ij_, we modulate the interaction strengths between different components. Specifically, a larger *λ*_ij_ value corresponds to stronger attractive interactions and higher hydrophobicity, while *λ*_ij_ = 0 leads to hydrophilic conditions.

We performed Langevin dynamics simulations using a friction coefficient *γ*_i_ = *m*_i_/*t*_damp_, where *m*_i_ is the mass of residues and *t*_damp_ = 1000 ps at a constant temperature of *T* = 300 K. Simulations were run for a total of 0.5 μs with a time step of 10 fs. All simulations were conducted using HOOMD-blue (version 2.9.3) with additional features implemented via the azplugins extension (version 0.10.1). The system was simulated in a cubic box (400 Å × 400 Å × 400 Å) containing 500 A and 500 B polymers, which together formed a stable droplet with radius of gyration 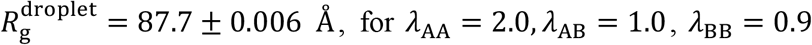, for *λ*_AA_ = 2.0, *λ*_AB_ = 1.0, *λ*_BB_ = 0.9. For reference, the radii of the pure A and B droplets were 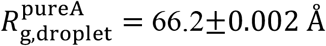 and 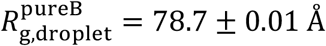, respectively, reflecting the higher hydrophobicity of the A species. Following the formation of the stable droplet, small molecules were then introduced by randomly placing additional C beads around the preformed droplet, corresponding to the target concentrations.

To quantify the effective interaction strength between the chains in the presence of small molecules, we first calculated the second virial coefficient, 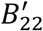, and then normalized it to obtain the dimensionless coefficient 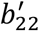. Specifically, 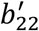 was obtained by dividing 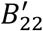 by its reference-state value computed in the absence of interchain attractions, thereby eliminating the influence of chain volume. 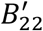 was calculated using the following integral:

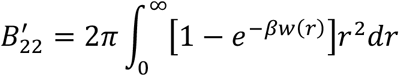

where *r* is the distance between the centers of mass (COM) of two chains, *w*(*r*) is the potential of mean force (PMF) between chains, and *β* = 1/*k*_B_ *T*. The PMF was obtained via umbrella sampling simulations performed in LAMMPS using the COLVARS module. Each simulation was conducted in a cubic box of edge length 200 Å containing a single A chain and a single B chain, along with an appropriate number of C particles according to the specified concentrations (25 mM, 50 mM, 100 mM, 200 mM and 500 mM). Umbrella sampling windows were spaced every 3 Å, covering the range from 0 to 81 Å. A harmonic bias potential with a spring constant of 0.2 kcal/(mol Å^2^: was applied in each window. Each window was simulated for 100 ns using a time step of 10 fs.

## Result and Discussion

### Morphological Transitions in Condensates Mediated by Interaction Strength with Small-Molecule Cosolutes

To establish a reference, we initially simulated the binary system consisting of A and B chains with the solvent modeled implicitly (**Fig. 1a**). We fixed the interactions between A-A monomer pairs to *λ*_AA_ = 2.0 and systematically varied the interaction strengths of B-B monomer pairs and A-B monomer pairs.

By adjusting the relative monomer interaction strengths between B-B and A-B, we observed distinct morphologies, including core-shell, dewetted, and mixed structures (**Fig. 1b**). These structural variations were further validated using concentration profiles, radial concentration distributions for spherical droplets and concentration gradients along the vector connecting the COM of the A-rich phase to that of the B-rich phase for non-spherical droplets (**Fig. 1c-d, Fig. S1**). When the A-B monomer interaction was weaker than or comparable to the B-B monomer interaction but still weaker than the A-A monomer interaction (*λ*_AB_ ≤ *λ*_BB_ < *λ*_AA_), the B chains preferentially interacted with themselves rather than with the A chains, leading to the formation of a dewetted structure, as indicated by the minimal overlap in the concentration profile along the horizontal axis (**Fig. 1c**). When *λ*_AB_ = *λ*_AA_, a mixed structure emerged (**Fig. 1b**) and when the monomer interaction between A-B was stronger than or comparable to the B-B pairs (*λ*_BB_ ≤ *λ*_AB_ < *λ*_AA_), a core-shell structure formed, wherein the A chains consistently occupied the inner region. This result aligns with previous findings that components with stronger homotypic interactions preferentially localize to the core^16,39^. Overall, these results highlight the pivotal role of homotypic and heterotypic interactions in dictating the morphology of multicomponent condensates.

Based on the results above, we selected *λ*_AB_ = 1.0 with *λ*_BB_ = 0.9 as the initial condition, which formed a stable core-shell structure. We then investigated the effect of introducing small molecules on the condensate’s morphology. The small molecule was modeled as a single bead (C), with only excluded volume interactions between C particles. Specifically, C particles were introduced at a concentration of 100 mM, initially positioned around the preformed core-shell droplet. Subsequently, the interactions of C with monomers A and B were systematically varied. To assess structural changes, we analyzed the COM distance between the phases formed by A and B chains 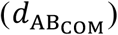, along with configuration snapshots (cross-sectional view). In a core-shell structure, the core-forming component A and the shell-forming component B occupy distinct spatial regions within the condensate with a small 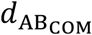. Conversely, a transition toward a dewetted morphology is indicated by an increase in 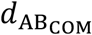, reflecting greater spatial segregation between A-rich and B-rich phases. For certain A-C and B-C interactions, we observed that the core-shell structure gradually transitioned into a dewetted morphology (**Fig. 2a(i),(ii),(iv),(v))**. When *λ*_AC_ = 0 was held constant, increasing *λ*_BC_ resulted in a greater partitioning ratio of C (defined as the fraction of C particles located inside the condensate, shown by green pentagons, right *y*-axis). To quantify the geometric relationship between A-rich and B-rich phases in dewetted morphologies, we calculated the intersection angle, defined in the schematic of **Fig. 2b** as the angle between vectors connecting the intersection point to the COM of each phase (see details in SI). Correspondingly, the intersection angle emerged and gradually increased from an initially undefined state, indicating a transition from a core-shell structure to dewetting, with the degree of dewetting increasing progressively (**Fig.2b**). When *λ*_BC_ = 0 was fixed and *λ*_AC_ was increased, more C particles entered the condensate, consistent with recent findings that particle partitioning into biomolecular condensates is governed by condensate-particle interaction strength, even though the condensates were covered by a layer of non-attractive B particles (*λ*_BC_ = 0)^41^. Interestingly, although the partition ratio increased with *λ*_AC_, a progressive dewetting trend was not observed. Instead, dewetting occurred abruptly and only at *λ*_AC_ = 3.0 (**Fig.2c**), suggesting that, in addition to the interaction strength with the small molecules, other factors also contribute to the observed morphological transitions, a point we will return to later.

**Fig. 2.**
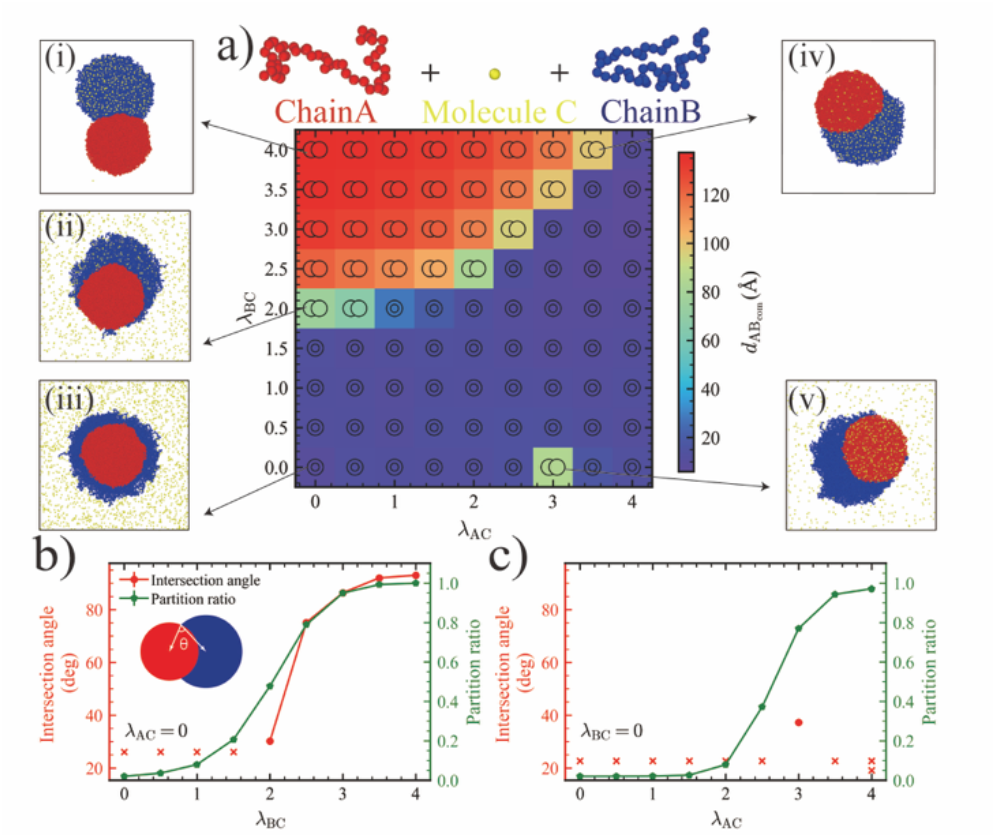
(a) Distance between the COM of the phases formed by the A and B chains, 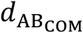, as a function of *λ*_BC_ and *λ*_AC_. Colors from purple to red indicate increasing 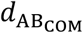. Representative cross-sectional snapshots (i-v) illustrate different regimes, with colors corresponding to the schematic component colors. (b) Intersection angle (red circles, left *y*-axis) and partition ratio of C particles (green pentagons, right *y*-axis) as functions of *λ*_BC_ at *λ*_AC_ = 0. The schematic illustrates the definition of the intersection angle. (c) Intersection angle (red circles, left *y*-axis) and partition ratio (green pentagons, right *y*-axis) as functions of *λ*_AC_ at *λ*_BC_ = 0. Cross symbols indicate cases where no intersection angle could be defined. Concentration of C equals to 100 mM.

A possible explanation for the observed morphological transitions is that the small molecules alter the effective interaction strengths among the macromolecular components of the condensate. To quantify these changes, we computed the normalized chain-chain second virial coefficient, 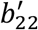, in the presence of C particles. A positive 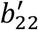 indicates effective repulsion, while negative 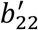 reflects effective attraction. Since A-A chain interactions are significantly stronger (with substantially more negative 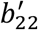) than other pairs, the A chains consistently form the inner phase in the core-shell morphologies. Therefore, to understand the morphological transition between core-shell and dewetted states, we focused primarily on the A-B and B-B chain interactions.

In the absence of attraction between small molecules and chains (*λ*_AC_ = 0, *λ*_BC_ = 0), the value of 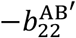 exceeds that of 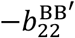, indicating that a B chain prefers to associate with an A chain over itself, which is consistent with the monomer-monomer interactions (*λ*_AB_ = 1.0 > *λ*_BB_ = 0.9), to gain more enthalpy, resulting in a core-shell structure. However, as *λ*_BC_ is increased (with *λ*_AC_ = 0; **Fig.3a**), 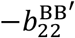 (green pentagons) increases significantly, reflecting stronger B-B interactions mediated by C particles, while 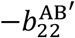 (red circles) decreases due to the purely repulsive A-C interactions. At 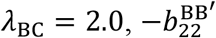 surpasses 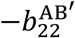, indicating that the B chains now favor homotypic interactions over heterotypic interactions, which is consistent with the dewetting observed in the condensate simulations. When *λ*_AC_ is increased at fixed *λ*_BC_ = 0 (**Fig. 3b**), the B-B chain interaction remains unchanged, while the A-B chain interactions weaken with increasing *λ*_AC_. As shown previously, when A-B chain interactions dominate over B-B, core-shell structures are favored. However, when *λ*_AC_ ≥ 3.0, the B-B chain interaction becomes effectively stronger. Based on these two-chain calculations, dewetting would be expected in the multi-chain condensate simulations; yet, we only observe dewetting at *λ*_AC_ = 3.0, suggesting a deviation from the expected trend (see discussion below).

**Fig. 3.**
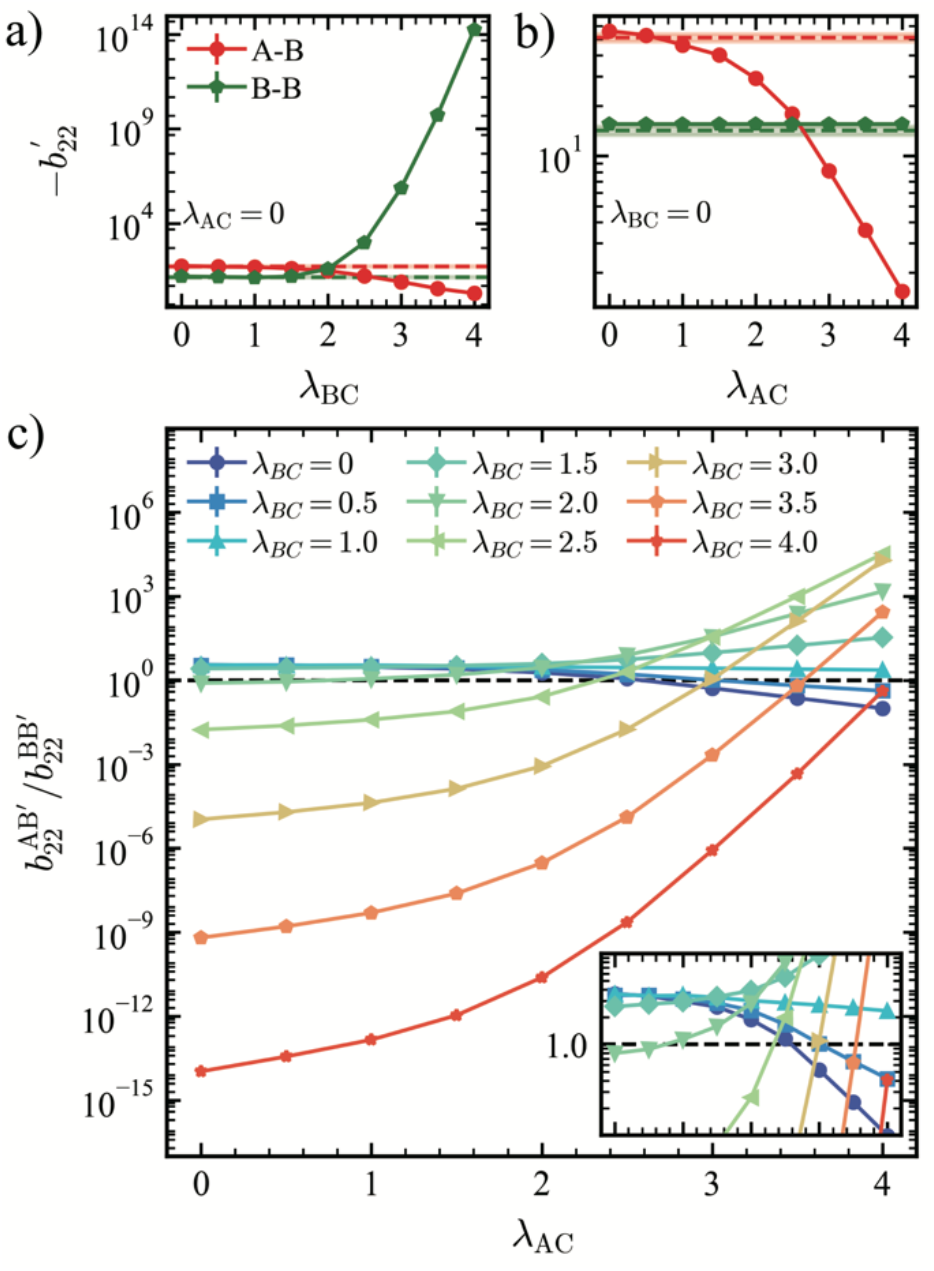
(a) Normalized chain-chain second virial coefficients 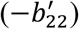 for chain A-chain B (red circles) and chain B-chain B (green pentagons) as functions of *λ*_BC_ at *λ*_AC_ = 0. Red and green horizontal dashed lines represent the corresponding −*b*_22_ values in the absence of C particles. The shaded regions around each dashed line indicate the associated error bars. (b) −*b*_22_ values for chain A-chain B (red circles) and chain B-chain B (green pentagons) as functions of *λ*_AC_ at *λ*_BC_ = 0. Horizontal dashed lines and shaded error regions are defined as in (a). (c) Ratio 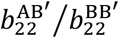 as a function of *λ* for various *λ*, with color from purple to red indicating increasing *λ*_BC_. The inset shows a zoomed-in view of the region around a ratio of 1, using the same x-axis range as the main plot.

Despite these differences, the ratio 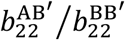 generally serves as a useful metric for capturing the relative strength of A-B versus B-B chain interactions and for explaining the observed morphological transitions (**Fig. 3c, Fig. S2**). When the ratio exceeds one, A-B chain interactions are stronger, leading to the core-shell structure to maximize heterotypic contacts. Conversely, when the ratio falls below one, B-B chain interactions dominate, driving B chains to self-associate, and resulting in dewetting. Additionally, a greater difference between 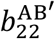 and 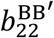 corresponds to a more pronounced dewetting behavior, consistent with changes in 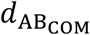.

### Stoichiometry-Dependent Modulation of Condensate Morphology by Small Molecules

While most of the observed condensate morphologies across various combinations of *λ*_AC_ and *λ*_BC_ show strong agreement with our two-chain theoretical calculations, a few notable discrepancies were identified. These inconsistencies primarily occurred at high values of *λ*_AC_ or *λ*_BC_ (≥ 3) and were associated with high partition ratios (**Fig. S3**), meaning that nearly all C particles were absorbed into the condensates.

We attribute these deviations to a mismatch between the dilute conditions for chains assumed in our theoretical calculations and the high-density environment of the condensate simulations. Although the concentration of the C particles was held constant across systems, the stoichiometric ratio between the small molecules and the macromolecules varied. Compared to our two-chain calculations, the condensate simulations involve a lower stoichiometric ratio, making fewer small molecules available to mediate effective interactions between chains. In condensate simulations with strong small molecule-macromolecule interactions, the C particles fully partition into the condensate, but their number is insufficient to substantially modulate the interaction among A and B chains. As a result, even in regimes where the effective interaction strengths determined from our two-chain calculations predict dewetting, the condensates exhibit a core-shell morphology instead. These observations suggest that stoichiometry, the ratio of small molecules to chains, plays a critical role in determining condensate morphology. Motivated by this insight, we systematically varied the concentration of C particles while keeping the concentrations of A and B chains constant.

We set *λ*_BC_ = 0 and systematically varied the interaction strength between C particles and A chains (*λ*_AC_) at different concentrations of C (**Fig. 4a**). When *λ*_AC_ = 0 and 1.0, the interaction between C and A monomers is weaker than the homotypic A-A monomer interaction (*λ*_AA_ = 2.0), and thus the condensates maintain their original core-shell morphology. However, when *λ*_AC_ ≥ 2.0, a clear concentration-dependent morphological transition is observed. At low concentrations of C, the condensates still retain their core-shell structure, but dewetting occurs as the concentration increases beyond a threshold. As the A-C attraction strength is increased further to *λ*_AC_ = 4.0, the condensate splits into orthogonal phases, one composed of B chains and one consisting of A chains and C molecules. To gain molecular-level insight into this behavior, we calculated the PMF between chain A-chain B and chain B-chain B for *λ*_AC_ = *λ*_AA_ = 2.0, which ensures that the A-A interactions remain less affected. As the concentration of component C increases, more C particles interact with chain A, effectively reducing the availability of A to interact with B. This competitive binding weakens the A-B attraction (**Fig. 4b**). Meanwhile, although less pronounced, B-B interactions are slightly enhanced due to the excluded volume effect of C. As a result, once the A-B chain interaction becomes weaker than the B-B chain interaction, dewetting is observed. These results demonstrate that the stoichiometry between small-molecule cosolutes and condensate can significantly influence the effective interactions among condensate components, thereby driving morphology changes.

**Fig. 4.**
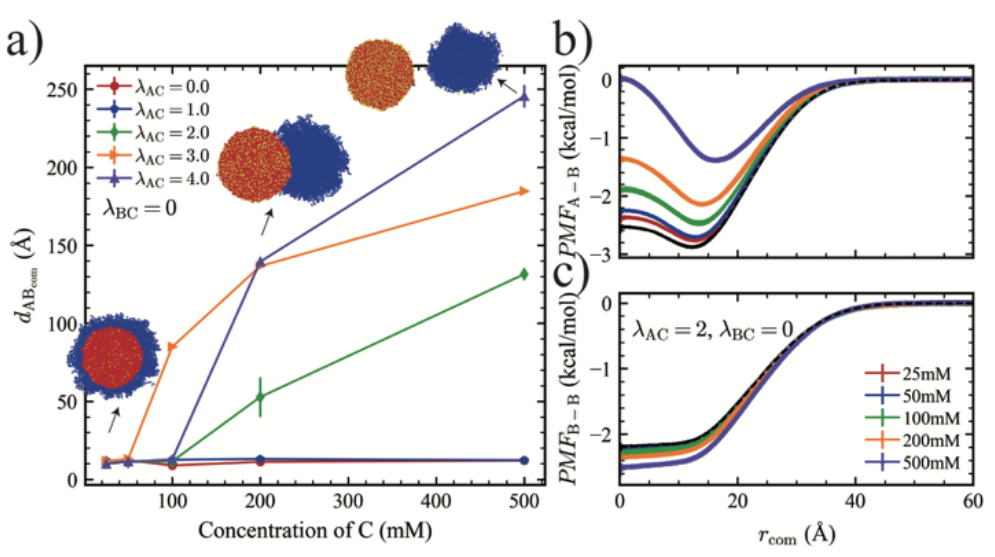
(a) 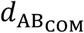 as a function of the concentration of component C under *λ*_BC_ = 0, with varying *λ*_AC_. Snapshots illustrate the condensate morphology for the condition *λ*_BC_ = 0, *λ*_AC_ = 4.0. (b-c) Potential of mean force (PMF) between (b) chain A and chain B, and between (c) chain B and chain B, as functions of the concentration of component C, for fixed *λ*_BC_ = 0, *λ*_AC_ = 2.0. The black dashed line indicates the PMF in the absence of particle C.

Based on the previous results, we further varied *λ*_BC_ while keeping *λ*_AC_ = 2.0 to systematically investigate how the interaction strength and stoichiometry between the condensate components and the small molecules jointly influence morphological transitions. At low concentration of C, consistent with earlier observations, increasing *λ*_BC_ initially had little effect on morphology; 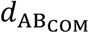remained relatively constant. However, beyond a threshold value, 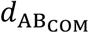 increased gradually, indicating a transition from a core-shell structure to a dewetted morphology (**Fig. 5a**). At higher C concentrations (200 mM and 500 mM), we observed a non-monotonic trend, a “re-dewetting” phenomenon, where dewetting initially appeared at low *λ*_BC_, then disappeared, and reemerged at higher *λ*_BC_ values (**Fig. 5a**). To understand this behavior, we analyzed the normalized second virial coefficients, 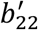, for both A-B and B-B chain pairs.

**Fig. 5.**
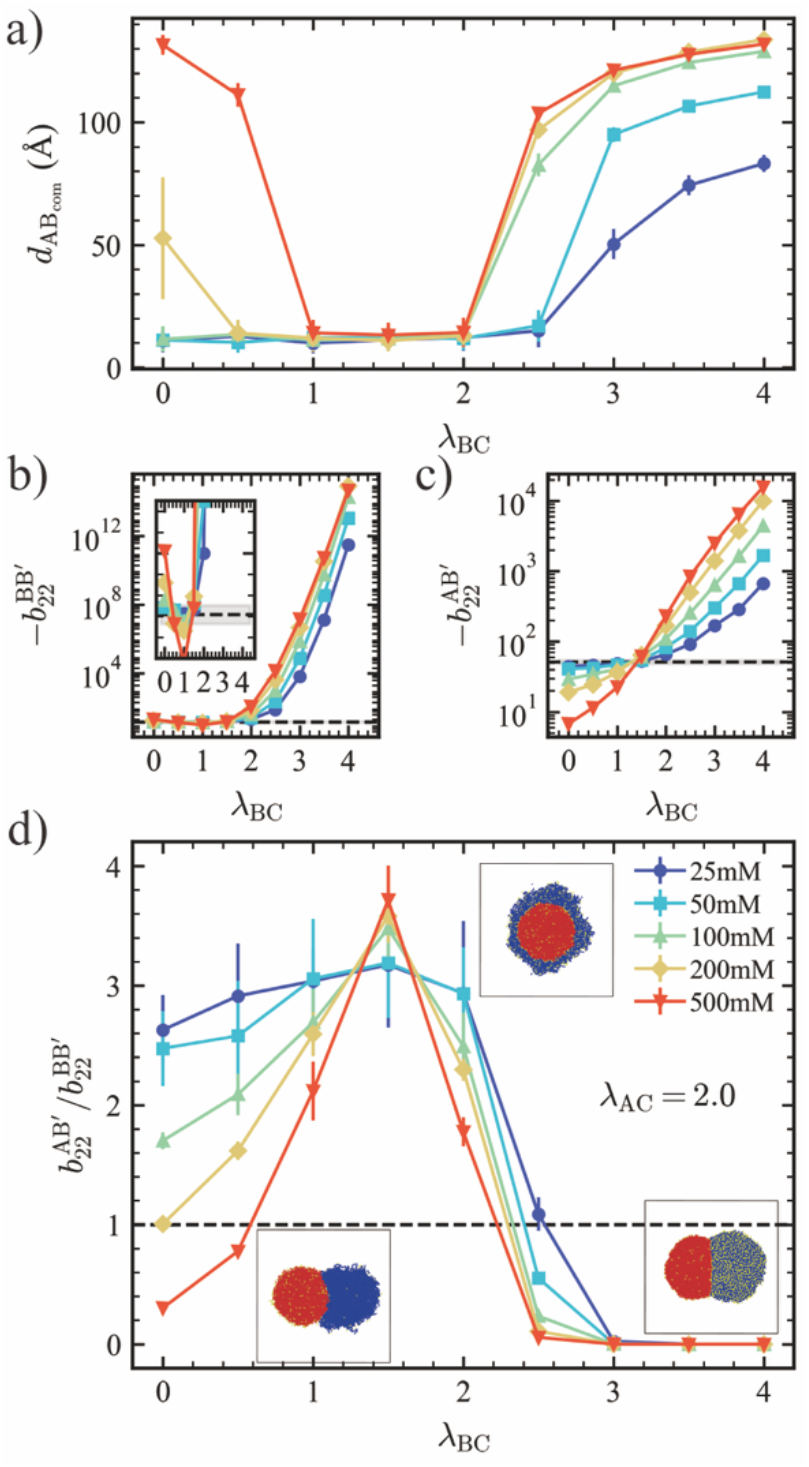
(a) 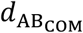 as a function of the *λ*_BC_ under different concentrations of C at *λ*_AC_ = 2.0. (b-c) 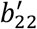 values for (b) chain B-chain B and (c) chain A-chain B as a function of *λ*_BC_ under different concentrations of C. Dashed lines represent the corresponding *b*_22_ in the absence of C. The inset in panel (b) provides a magnified view of values near the dashed line. (d) Ratio 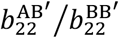 as a function of *λ*_BC_ for various concentrations of C. Snapshots illustrate the condensate morphologies at C = 500 mM for *λ*_BC_ = 0,1.5,4.0, respectively. All plots use the same color and marker legend as shown in panel (d).

When *λ*_BC_ = 0, due to volume exclusion between C-C and B-C pairs, the C particles occupy space, slightly enhancing B-B chain interactions compared to the no-C condition. As *λ*_BC_ increases, the effective B-B chain interaction initially decreases slightly (**Fig. 5b, inset**), which can be attributed to the relatively weak B-C attraction leading to the weaker volume exclusion effect and disrupting B-B interaction. However, upon further increasing *λ*_BC_, the stronger B-C interactions begin to promote attraction between B chains. This trend becomes more pronounced at higher concentrations of C. Although we did not directly modify A-B interactions, the presence of C also affects the effective interaction between chain A and chain B as a mediator. At *λ*_BC_ = 0, the A-B chain interaction is weaker than in the absence of C (**Fig. 5c**), because a portion of A interacts with C (*λ*_AC_ = 2.0), which is stronger than the A-B interaction. This competition weakens A-B interactions, and the effect is more pronounced at higher concentrations of C. However, as *λ*_BC_ increases, C can now interact with both A and B, indirectly enhancing the effective A-B interaction. This compensation effect becomes stronger as *λ*_BC_ increases further. Eventually, the effective A-B chain interaction exceeds the value observed in the absence of C, as more C particles bridge A and B interactions.

We continue to use the ratio 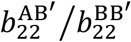 as a criterion which can be either greater or less than 1, corresponding to core-shell and dewetted morphologies, respectively (**Fig. 5d**). In the first dewetting regime at low *λ*_BC_, a high concentration of C enhances B-B chain interactions while weakening A-B chain interactions relative to the no-C system. Once the attraction between B-B pairs is stronger than for A-B pairs, dewetting occurs. The degree of this transition depends strongly on the concentration of C. As *λ*_BC_ continues to increase, the enhancement of A-B chain interactions becomes more pronounced than that of B-B, raising the ratio 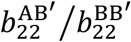 above 1 and resulting in a core-shell morphology. At even higher *λ*_BC_, B-B chain attraction once again surpasses that of A-B pairs, causing the reemergence of dewetting-a “second dewetting”.

Overall, our investigations demonstrate that the small-molecule cosolute modulates condensate morphology through its interactions with the constituent chains. This regulation is governed by both the interaction strength and stoichiometry between the small molecule and condensate components, which together determine the morphology of the condensate.

### Small Molecule-Induced Morphological Transitions in Miscible Multicomponent Condensates

As demonstrated in the previous sections, the morphology of multiphase condensates can be influenced by both the interaction strength and stoichiometry with the small molecules. However, many biologically relevant condensates are multicomponent yet exhibit a single-phase (mixed) morphology. Therefore, we investigated the impact of small molecules on fully mixed condensates by fixing *λ*_AA_ = *λ*_AB_ = *λ*_BB_ = 1.0 at the 100 mM concentration of C particles. We then systematically varied the interaction strengths between C and A, B monomers, and observed the emergence of distinct morphologies (**Fig. 6, Fig. S4**). Specifically, when *λ*_AC_ = 3.0, increasing *λ*_BC_ led to a gradual decrease in 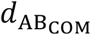. Combined with the corresponding snapshots, this trend revealed a sequence of morphological transitions: from a dewetted structure, to a core-shell structure with A chains occupying the core, to a fully mixed state, and finally to an inverse core-shell structure with B chains occupying the core (**Fig. 6a**). To explain these transitions, we again employed the 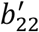 as a metric for the effective chain-chain interaction strengths (**Fig. 6b, Fig S5**). Unlike previous cases, the A-A interaction is no longer dominant, so both 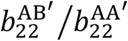 and 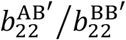 must be considered. When *λ* is small, both ratios are less than 1, indicating that A-B chain interactions are weaker than the corresponding homotypic interactions (i.e., 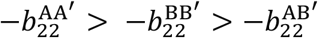). As a result, a dewetted structure is observed. As *λ* increases, the ratio 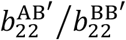 surpasses 1, while 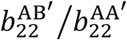 remains below 1, which implies that A-B chain interactions become stronger than B-B, while A-A interactions remain dominant, leading to the formation of a core-shell structure with A chains occupying the inner phase. Notably, at *λ*_BC_ = 1.5, we observe a discrepancy between the condensate simulations and the theoretical prediction based on the effective chain-chain interactions, which we attribute to stoichiometry effects, as previously discussed. Upon further increasing *λ*_BC_, all three interaction strengths approach equivalence, i.e., 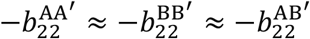, resulting in a fully mixed structure. If *λ* continues to increase, B-B chain interactions eventually become the strongest, leading to the formation of an inverse core-shell structure with B chains occupying the interior. These results demonstrate that in a single-phase multicomponent system, the small molecules can effectively regulate condensate morphology by altering the relative effective interaction strengths among the constituent chains. **Conclusions**

**Fig. 6.**
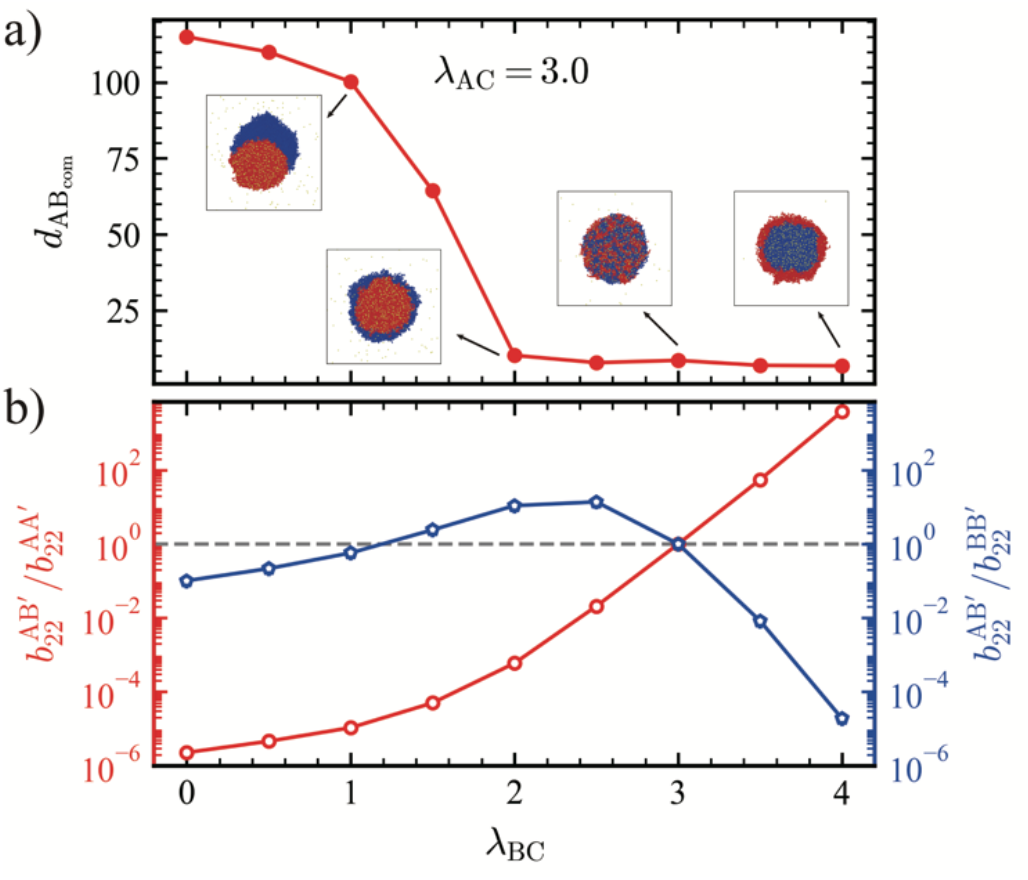
(a) 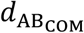 as a function of *λ*_BC_ at fixed *λ*_AC_ = 3.0. Snapshots show cross-sectional views of representative condensate morphologies. (b) Ratio 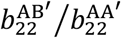 (left *y*-axis, red) and ratio 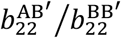 (right *y*-axis, blue) as a function of *λ*_BC_ under *λ*_AC_ = 3.0.

In this study, we investigated how small-molecule cosolutes modulate the morphology of multicomponent condensates from a molecular interaction perspective. By systematically varying the interaction strengths between the small molecules and each condensate component, we demonstrated that small molecules can effectively reshape the condensate morphology. To interpret these transitions, we computed normalized effective second virial coefficients,, to quantify homotypic and heterotypic interactions between two chains in the presence of small molecules, and demonstrated that the morphological changes arise from shifts in their relative strengths.

By systematically varying the concentration of the small molecules, we further showed that concentration-dependent modulation of effective interactions can account for highly non-linear morphological transitions observed in simulations. We also extended our analysis to multicomponent condensates that are fully miscible in the absence of cosolutes, and found that tuning the relative interaction strengths through small molecules could induce transitions to different morphologies such as dewetted, core-shell, or inverse core-shell.

This molecular interaction-based framework can also be extended to macromolecular regulators, such as RNAs and proteins, which have similarly been shown to influence condensate morphology^18,19^. Although these macromolecules often exhibit multivalency, the resulting morphological transitions in multicomponent condensates still arise from the same fundamental principle, the balance between homotypic and heterotypic interactions. Whether mediated by small-molecule cosolutes or larger biomolecules such as proteins and nucleic acids, modulation of these interactions governs the structural organization of condensates. Together, our findings provide a mechanistic basis for understanding and engineering how condensate morphology can be tuned through interaction-driven processes, offering insights that may inform future experimental and computational strategies to control condensate structure and function across diverse biological contexts.

## Supporting Information

Analysis methods, data for density profile, normalized second virial coefficients between components under different conditions, partition ratio diagram and state diagram.

## Acknowledgements

This material is based on the work supported by the National Institute of General Medical Science of the National Institutes of Health under grant R35GM153388. A.N. acknowledges funding by the Deutsche Forschungsgemeinschaft (DFG, German Research Foundation) through Project 470113688. Y.C.K. is supported by the Office of Naval Research via the U.S. Naval Research base program. We gratefully acknowledge the computational resources provided by the Texas A&M High Performance Research Computing (HPRC) to complete this work.

## Reference

(1) Wei, M.; Wang, X.; Qiao, Y. Multiphase Coacervates: Mimicking Complex Cellular Structures through Liquid–Liquid Phase Separation. Chemical Communications 2024, 60 (90), 13169–13178. 10.1039/D4CC04533E.

(2) Banani, S. F.; Lee, H. O.; Hyman, A. A.; Rosen, M. K. Biomolecular Condensates: Organizers of Cellular Biochemistry. Nat Rev Mol Cell Biol 2017, 18 (5), 285–298. 10.1038/nrm.2017.7.

(3) Fare, C. M.; Villani, A.; Drake, L. E.; Shorter, J. Higher-Order Organization of Biomolecular Condensates. Open Biology 2021, 11 (6), 210137. 10.1098/rsob.210137.

(4) Feric, M.; Vaidya, N.; Harmon, T. S.; Mitrea, D. M.; Zhu, L.; Richardson, T. M.; Kriwacki, R. W.; Pappu, R. V.; Brangwynne, C. P. Coexisting Liquid Phases Underlie Nucleolar Subcompartments. Cell 2016, 165 (7), 1686–1697. 10.1016/j.cell.2016.04.047.

(5) Jain, S.; Wheeler, J. R.; Walters, R. W.; Agrawal, A.; Barsic, A.; Parker, R. ATPase-Modulated Stress Granules Contain a Diverse Proteome and Substructure. Cell 2016, 164 (3), 487–498. 10.1016/j.cell.2015.12.038.

(6) Lafontaine, D. L. J.; Riback, J. A.; Bascetin, R.; Brangwynne, C. P. The Nucleolus as a Multiphase Liquid Condensate. Nat Rev Mol Cell Biol 2021, 22 (3), 165–182. 10.1038/s41580-020-0272-6.

(7) Fei, J.; Jadaliha, M.; Harmon, T. S.; Li, I. T. S.; Hua, B.; Hao, Q.; Holehouse, A. S.; Reyer, M.; Sun, Q.; Freier, S. M.; Pappu, R. V.; Prasanth, K. V.; Ha, T. Quantitative Analysis of Multilayer Organization of Proteins and RNA in Nuclear Speckles at Super Resolution. Journal of Cell Science 2017, 130 (24), 4180–4192. 10.1242/jcs.206854.

(8) Wheeler, J. R.; Matheny, T.; Jain, S.; Abrisch, R.; Parker, R. Distinct Stages in Stress Granule Assembly and Disassembly. eLife 2016, 5, e18413. 10.7554/eLife.18413.

(9) Boeynaems, S.; Holehouse, A. S.; Weinhardt, V.; Kovacs, D.; Van Lindt, J.; Larabell, C.; Van Den Bosch, L.; Das, R.; Tompa, P. S.; Pappu, R. V.; Gitler, A. D. Spontaneous Driving Forces Give Rise to protein–RNA Condensates with Coexisting Phases and Complex Material Properties. Proceedings of the National Academy of Sciences 2019, 116 (16), 7889–7898. 10.1073/pnas.1821038116.

(10) Boisvert, F.-M.; van Koningsbruggen, S.; Navascués, J.; Lamond, A. I. The Multifunctional Nucleolus. Nat Rev Mol Cell Biol 2007, 8 (7), 574–585. 10.1038/nrm2184.

(11) Mao, S.; Chakraverti-Wuerthwein, M. S.; Gaudio, H.; Košmrlj, A. Designing the Morphology of Separated Phases in Multicomponent Liquid Mixtures. Phys. Rev. Lett. 2020, 125 (21), 218003. 10.1103/PhysRevLett.125.218003.

(12) Pyo, A. G. T.; Zhang, Y.; Wingreen, N. S. Surface Tension and Super-Stoichiometric Surface Enrichment in Two-Component Biomolecular Condensates. iScience 2022, 25 (2), 103852. 10.1016/j.isci.2022.103852.

(13) Gouveia, B.; Kim, Y.; Shaevitz, J. W.; Petry, S.; Stone, H. A.; Brangwynne, C. P. Capillary Forces Generated by Biomolecular Condensates. Nature 2022, 609 (7926), 255–264. 10.1038/s41586-022-05138-6.

(14) Ye, S.; Latham, A. P.; Tang, Y.; Hsiung, C.-H.; Chen, J.; Luo, F.; Liu, Y.; Zhang, B.; Zhang, X. Micropolarity Governs the Structural Organization of Biomolecular Condensates. bioRxiv March 30, 2023, p 2023.03.30.534881. 10.1101/2023.03.30.534881.

(15) Welles, R. M.; Sojitra, K. A.; Garabedian, M. V.; Xia, B.; Wang, W.; Guan, M.; Regy, R. M.; Gallagher, E. R.; Hammer, D. A.; Mittal, J.; Good, M. C. Determinants That Enable Disordered Protein Assembly into Discrete Condensed Phases. Nat. Chem. 2024, 16 (7), 1062–1072. 10.1038/s41557-023-01423-7.

(16) Rana, U.; Xu, K.; Narayanan, A.; Walls, M. T.; Panagiotopoulos, A. Z.; Avalos, J. L.; Brangwynne, C. P. Asymmetric Oligomerization State and Sequence Patterning Can Tune Multiphase Condensate Miscibility. Nat. Chem. 2024, 16 (7), 1073–1082. 10.1038/s41557-024-01456-6.

(17) Zhorabek, F.; Sandupama Abesekara, M.; Liu, J.; Dai, X.; Huang, J.; Chau, Y. Construction of Multiphasic Membraneless Organelles towards Spontaneous Spatial Segregation and Directional Flow of Biochemical Reactions. Chemical Science 2023, 14 (4), 801–811. 10.1039/D2SC05438H.

(18) Kaur, T.; Raju, M.; Alshareedah, I.; Davis, R. B.; Potoyan, D. A.; Banerjee, P. R. Sequence-Encoded and Composition-Dependent Protein-RNA Interactions Control Multiphasic Condensate Morphologies. Nat Commun 2021, 12 (1), 872. 10.1038/s41467-021-21089-4.

(19) Rai, S. K.; Khanna, R.; Avni, A.; Mukhopadhyay, S. Heterotypic Electrostatic Interactions Control Complex Phase Separation of Tau and Prion into Multiphasic Condensates and Co-Aggregates. Proceedings of the National Academy of Sciences 2023, 120 (2), e2216338120. 10.1073/pnas.2216338120.

(20) Bai, Q.; Chen, X.; Chen, J.; Liu, Z.; Lin, Y.; Yang, S.; Liang, D. Morphology and Dynamics of Coexisting Phases in Coacervate Solely Controlled by Crowded Environment. ACS Macro Lett. 2022, 11 (9), 1107–1111. 10.1021/acsmacrolett.2c00409.

(21) Bai, Q.; Liu, Z.; Chen, J.; Liang, D. Crowded Environment Regulates the Coacervation of Biopolymers via Nonspecific Interactions. Biomacromolecules 2023, 24 (1), 283–293. 10.1021/acs.biomac.2c01129.

(22) Love, C.; Steinkühler, J.; Gonzales, D. T.; Yandrapalli, N.; Robinson, T.; Dimova, R.; Tang, T.-Y. D. Reversible pH-Responsive Coacervate Formation in Lipid Vesicles Activates Dormant Enzymatic Reactions. Angewandte Chemie International Edition 2020, 59 (15), 5950–5957. 10.1002/anie.201914893.

(23) de Haas, R. J.; Ganar, K. A.; Deshpande, S.; de Vries, R. pH-Responsive Elastin-Like Polypeptide Designer Condensates. ACS Appl. Mater. Interfaces 2023, 15 (38), 45336–45344. 10.1021/acsami.3c11314.

(24) Lu, T.; Spruijt, E. Multiphase Complex Coacervate Droplets. J. Am. Chem. Soc. 2020, 142 (6), 2905–2914. 10.1021/jacs.9b11468.

(25) Chen, X.; Chen, E.-Q.; Yang, S. Multiphase Coacervation of Polyelectrolytes Driven by Asymmetry of Charged Sequence. Macromolecules 2023, 56 (1), 3–14. 10.1021/acs.macromol.2c01205.

(26) Zhu, L.; Pan, Y.; Hua, Z.; Liu, Y.; Zhang, X. Ionic Effect on the Microenvironment of Biomolecular Condensates. J. Am. Chem. Soc. 2024, 146 (20), 14307–14317. 10.1021/jacs.4c04036.

(27) Hautke, A.; Ebbinghaus, S. The Emerging Role of ATP as a Cosolute for Biomolecular Processes. Biological Chemistry 2023, 404 (10), 897–908. 10.1515/hsz-2023-0202.

(28) Li, S.; Wang, Y.; Lai, L. Small Molecules in Regulating Protein Phase Separation. Acta Biochim Biophys Sin (Shanghai) 2023, 55 (7), 1075–1083. 10.3724/abbs.2023106.

(29) Babinchak, W. M.; Dumm, B. K.; Venus, S.; Boyko, S.; Putnam, A. A.; Jankowsky, E.; Surewicz, W. K. Small Molecules as Potent Biphasic Modulators of Protein Liquid-Liquid Phase Separation. Nat Commun 2020, 11 (1), 5574. 10.1038/s41467-020-19211-z.

(30) Kang, J.; Lim, L.; Song, J. ATP Enhances at Low Concentrations but Dissolves at High Concentrations Liquid-Liquid Phase Separation (LLPS) of ALS/FTD-Causing FUS. Biochemical and Biophysical Research Communications 2018, 504 (2), 545–551. 10.1016/j.bbrc.2018.09.014.

(31) Patel, A.; Malinovska, L.; Saha, S.; Wang, J.; Alberti, S.; Krishnan, Y.; Hyman, A. A. ATP as a Biological Hydrotrope. Science 2017, 356 (6339), 753–756. 10.1126/science.aaf6846.

(32) Hayes, M. H.; Peuchen, E. H.; Dovichi, N. J.; Weeks, D. L. Dual Roles for ATP in the Regulation of Phase Separated Protein Aggregates in Xenopus Oocyte Nucleoli. eLife 2018, 7, e35224. 10.7554/eLife.35224.

(33) Wang, Z.; Chen, D.; Guan, D.; Liang, X.; Xue, J.; Zhao, H.; Song, G.; Lou, J.; He, Y.; Zhang, H. Material Properties of Phase-Separated TFEB Condensates Regulate the Autophagy-Lysosome Pathway. Journal of Cell Biology 2022, 221 (5), e202112024. 10.1083/jcb.202112024.

(34) Li, P.; Banjade, S.; Cheng, H.-C.; Kim, S.; Chen, B.; Guo, L.; Llaguno, M.; Hollingsworth, J. V.; King, D. S.; Banani, S. F.; Russo, P. S.; Jiang, Q.-X.; Nixon, B. T.; Rosen, M. K. Phase Transitions in the Assembly of Multivalent Signalling Proteins. Nature 2012, 483 (7389), 336–340. 10.1038/nature10879.

(35) Han, T. W.; Kato, M.; Xie, S.; Wu, L. C.; Mirzaei, H.; Pei, J.; Chen, M.; Xie, Y.; Allen, J.; Xiao, G.; McKnight, S. L. Cell-Free Formation of RNA Granules: Bound RNAs Identify Features and Components of Cellular Assemblies. Cell 2012, 149 (4), 768–779. 10.1016/j.cell.2012.04.016.

(36) Lyon, A. S.; Peeples, W. B.; Rosen, M. K. A Framework for Understanding the Functions of Biomolecular Condensates across Scales. Nat Rev Mol Cell Biol 2021, 22 (3), 215–235. 10.1038/s41580-020-00303-z.

(37) Cai, C.; Li, Y.; Lin, J.; Wang, L.; Lin, S.; Wang, X.-S.; Jiang, T. Simulation-Assisted Self-Assembly of Multicomponent Polymers into Hierarchical Assemblies with Varied Morphologies. Angewandte Chemie 2013, 125 (30), 7886–7890. 10.1002/ange.201210024.

(38) Chen, Z.; Huo, J.; Hao, L.; Zhou, J. Multiscale Modeling and Simulations of Responsive Polymers. Current Opinion in Chemical Engineering 2019, 23, 21–33. 10.1016/j.coche.2019.02.004.

(39) Li, N.; Nikoubashman, A.; Panagiotopoulos, A. Z. Self-Assembly of Polymer Blends and Nanoparticles through Rapid Solvent Exchange. Langmuir 2019, 35 (10), 3780–3789. 10.1021/acs.langmuir.8b04197.

(40) Rekhi, S.; Sundaravadivelu Devarajan, D.; Howard, M. P.; Kim, Y. C.; Nikoubashman, A.; Mittal, J. Role of Strong Localized vs Weak Distributed Interactions in Disordered Protein Phase Separation. J. Phys. Chem. B 2023, 127 (17), 3829–3838. 10.1021/acs.jpcb.3c00830.

(41) Kelley, F. M.; Ani, A.; Pinlac, E. G.; Linders, B.; Favetta, B.; Barai, M.; Ma, Y.; Singh, A.; Dignon, G. L.; Gu, Y.; Schuster, B. S. Controlled and Orthogonal Partitioning of Large Particles into Biomolecular Condensates. July 16, 2024. 10.1101/2024.07.11.603072.

