## Supplemental information for "Controlling Multicomponent Condensate Morphology via Additive-Modulated Interactions"

### Analysis Methods

#### Intersection Angle Calculation

To quantify the geometric relationship between A-rich and B-rich phases in dewetted morphologies, we first defined the axis connecting their respective centers of mass (COMs) as the y-axis. An orthogonal direction was designated as the x-axis, forming an x-y plane onto which all particle coordinates were projected. This plane was discretized into a $200 \times 200$ grid. For each simulation frame, number density maps were generated on this grid, and a contour line was extracted at 90% of the maximum density to represent the phase boundary. Each contour was fitted to a circle, and the intersection points of the two circles were identified. The intersection angle was then defined as the angle between the lines connecting each COM to the intersection point.

#### Density Profile Along the Horizontal Axis

To evaluate the spatial distribution of components in dewetted morphologies, we used the vector connecting the COMs of the A-rich and B-rich phases as the horizontal axis. The system was divided into 50 bins along this direction. The concentrations of A and B monomers were calculated in each bin to construct density profiles.

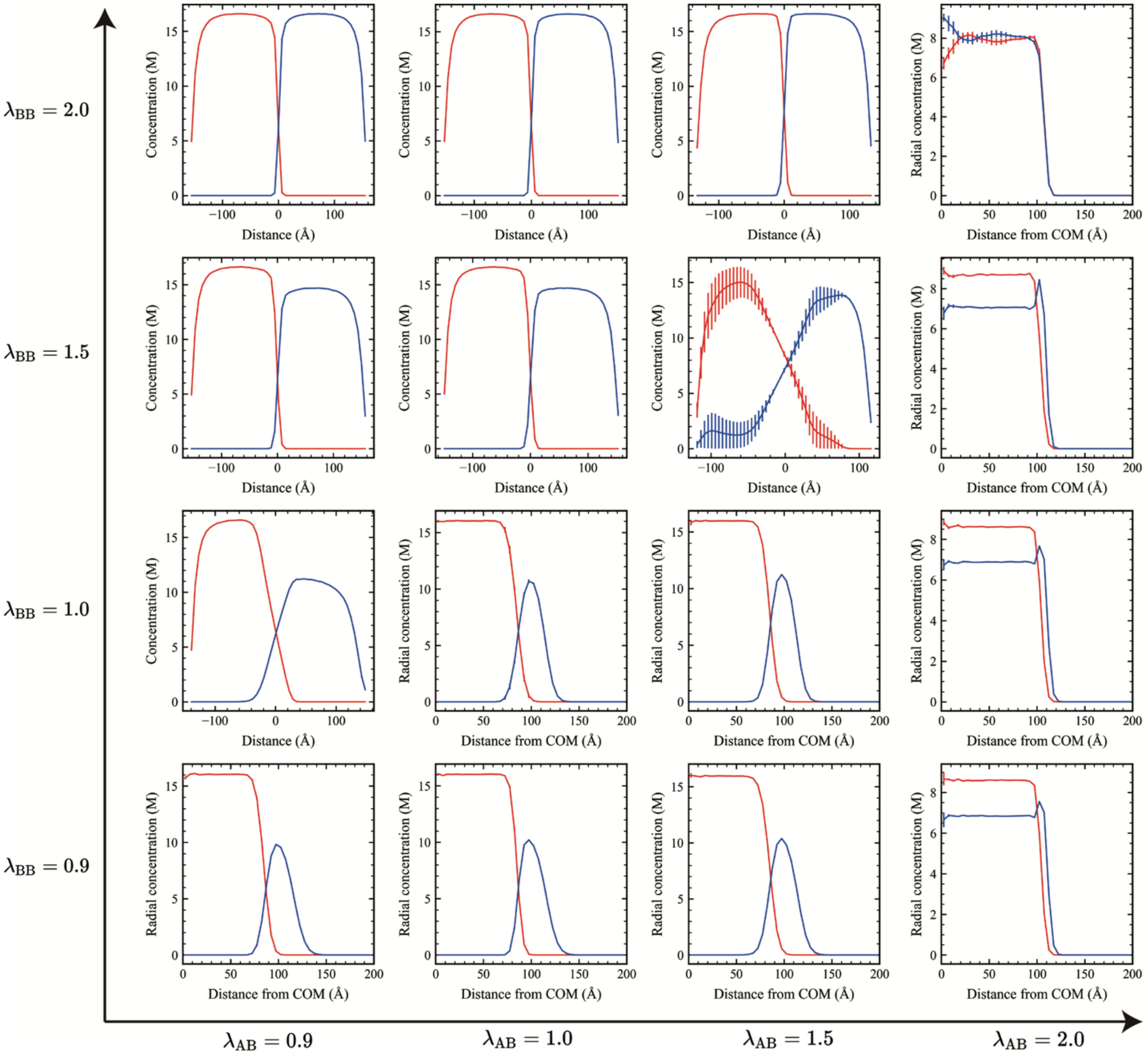


**Fig.S1** (a) Concentration profiles of A (red line) and B (blue line) monomers under different interaction strengths. Radial concentration distributions for spherical droplets and concentration gradients along the vector connecting the center-of-mass (COM) of the A-rich phase to that of the B-rich phase for non-spherical droplets.

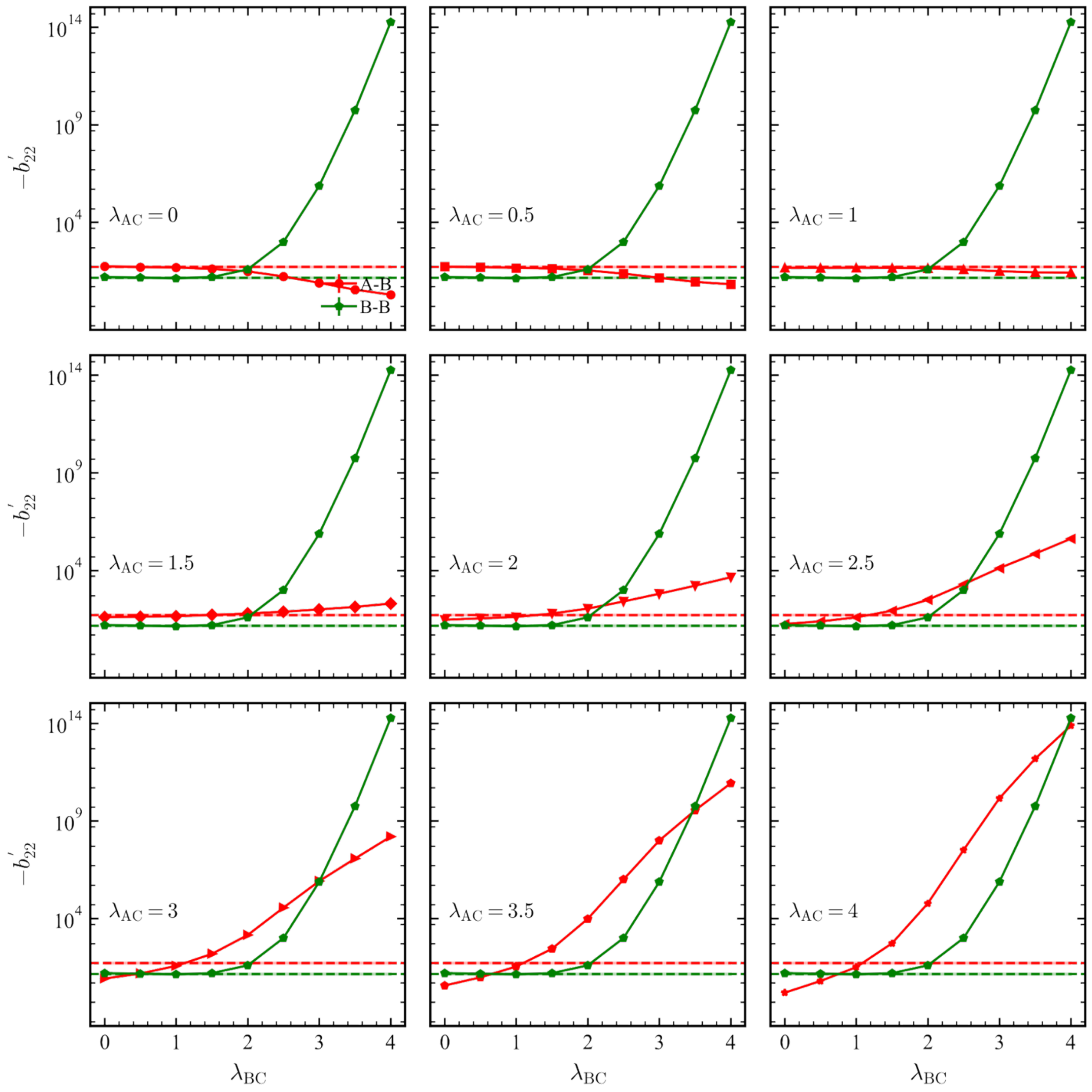


**Fig.S2** $-b_{22}$ values for Chain A-ChainB (red) and Chain B-Chain B(green) as functions of $\lambda_{\mathrm{BC}}$ at varies constant $\lambda_{\mathrm{AC}}$. Red and green horizontal dashed lines represent the corresponding $-b_{22}$ values in the absence of C particles. The shaded regions around each dashed line indicate the associated error bars.

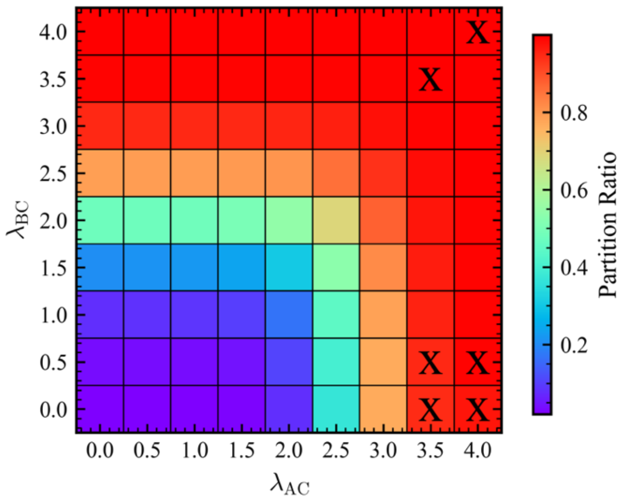


**Fig.S3** Partition ratio as a function of $\lambda_{BC}$ and $\lambda_{AC}$. Colors from purple to red indicate increasing Partition ratios. Squares marked with an "X" represent conditions where simulation results are inconsistent with theoretical predictions.

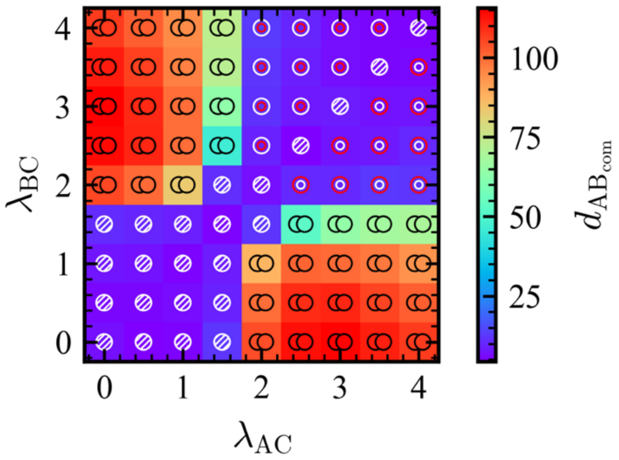


**Fig.S4** (a) $d_{AB_{COM}}$, as a function of $\lambda_{BC}$ and $\lambda_{AC}$. Colors from purple to red indicate increasing $d_{AB_{COM}}$. The marker on each colored square denotes the observed morphology: miscible (circle with hatch), dewetting (two connected circles), core-shell (concentric circles with red core and white shell), and inverse core–shell (concentric circles with white core and red shell).

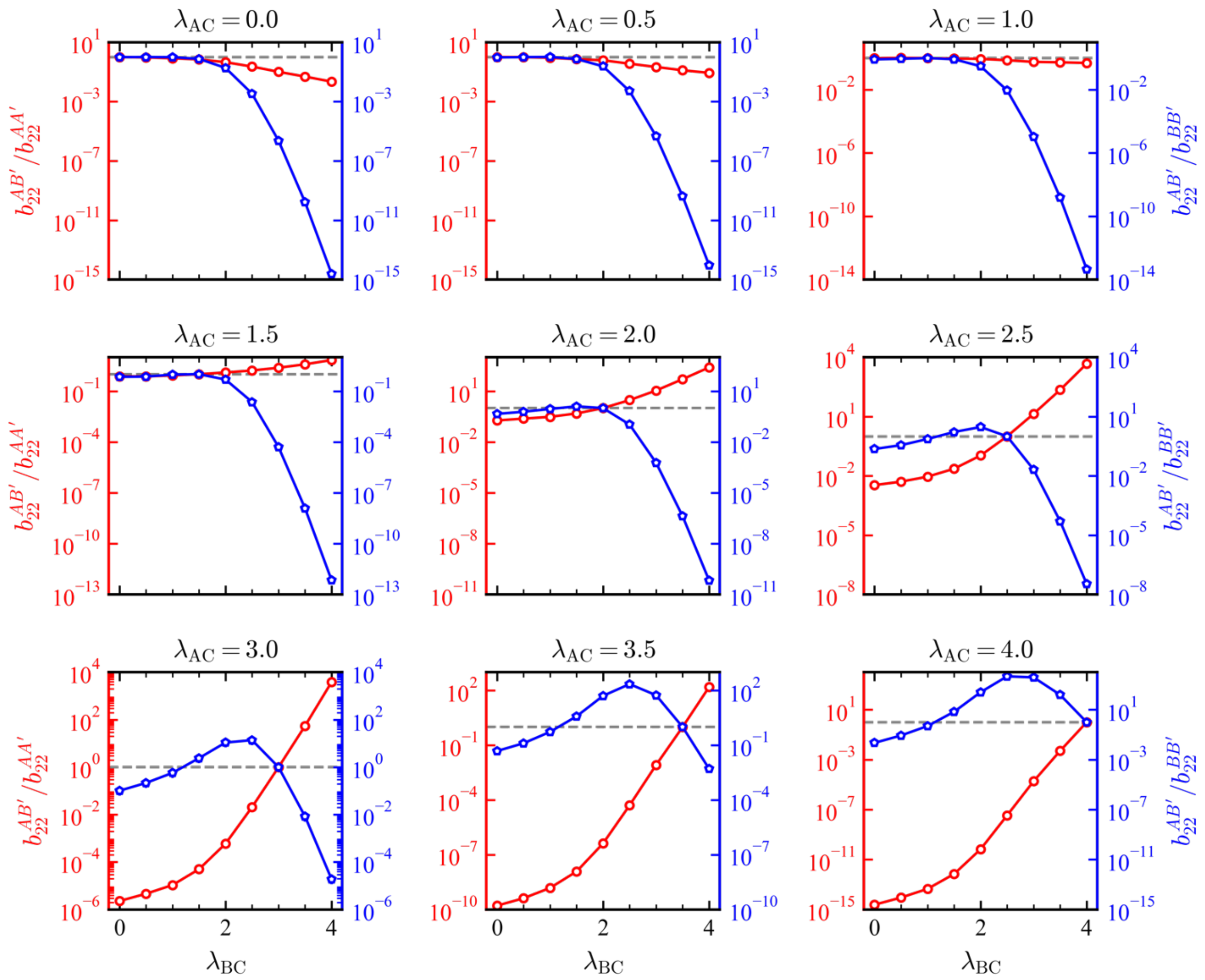

**Fig. S5** Ratio $b_{22}^{\mathrm{AB}'}/b_{22}^{\mathrm{AA}'}$ (left $y$-axis, red) and ratio $b_{22}^{\mathrm{AB}'}/b_{22}^{\mathrm{BB}'}$ (right $y$-axis, blue) as a function of $\lambda_{\mathrm{BC}}$ under different $\lambda_{\mathrm{AC}}$.
